# TCR-MHC Interaction Strength Defines Trafficking and Resident Memory Status of CD8 T cells in the Brain

**DOI:** 10.1101/263152

**Authors:** Anna Sanecka, Nagisa Yoshida, Elizabeth Motunrayo Kolawole, Harshil Patel, Brian D. Evavold, Eva-Maria Frickel

## Abstract

T cell receptor-Major histocompatibility complex (TCR-MHC) affinities span a wide range in a polyclonal T cell response, yet it is undefined how affinity shapes long-term properties of CD8 T cells during chronic infection with persistent antigen. Here, we investigate how the affinity of the TCR-MHC interaction shapes the phenotype of memory CD8 T cells in the chronically *Toxoplasma gondii-infected* brain. We employed CD8 T cells from three lines of transnuclear (TN) mice that harbour in their endogenous loci different T cell receptors specific for the same *Toxoplasma* antigenic epitope ROP7. The three TN CD8 T cell clones span a wide range of affinities to MHCI-ROP7. These three CD8 T cell clones have a distinct and fixed hierarchy in terms of effector function in response to the antigen measured as proliferation capacity, trafficking, T cell maintenance and memory formation. In particular, the T cell clone of lowest affinity does not home to the brain. The two higher affinity T cell clones show differences in establishing resident memory populations (CD103^+^) in the brain with the higher affinity clone persisting longer in the host during chronic infection. Transcriptional profiling of naïve and activated ROP7-specific CD8 T cells revealed that *Klf2* encoding a transcription factor that is known to be a negative marker for T cell trafficking is upregulated in the activated lowest affinity ROP7 clone. Our data thus suggest that TCR-MHC affinity dictates memory CD8 T cell fate at the site of infection.

## Introduction

CD8 T cells are a cornerstone of the adaptive immune defence to intracellular pathogens with their capacity to operate as antigen-experienced effector and memory cells. Pathogen-specific CD8 effector T cells rapidly expand and differentiate during the acute infection, followed by a phase of contraction and development of long-lived memory T cells (1,2). Most of our understanding of T cell responses to chronic infections is derived from models where pathogen control is incomplete and T cell become functionally impaired or exhausted over time (3,4). We thus lack knowledge of what drives long-lasting control of chronically persistent pathogens.

The interaction of the TCR with the pathogen antigenic epitope loaded on the MHC is essential in maintaining effective CD8 T cell control of persistent intracellular pathogens. The aβ TCR stochastically assembles and is selected during thymic development, and it is via this receptor that the immune system tunes the breath and strength of its response (2,5). Efforts have been made to elicit the effect of TCR-MHC affinity on the fate of the resulting T cells, however, often this relied on varying the antigenic peptide rather the TCR (2,6). The simple question of how T cells of different affinity to a given antigen fare during chronic infection remains unresolved.

In order to model a persistent chronic infection, we deemed a resistant mouse strain infected with the protozoan parasite *Toxoplasma gondii* to be most suitable. *Toxoplasma* is the most common parasitic infection in man, whereby in immunocompetent hosts the acute phase of infection is generally asymptomatic and proceeds to the chronic phase, which is incurable and defined by tissue cyst formation preferably in the brain. The parasite poses a serious health threat to immunocompromised individuals, especially AIDS patients. It is unclear how *Toxoplasma* maintains the intricate balance between survival and host defence. CD8 T cells and their ability to produce IFNγ have been shown to secure the latency of the parasitic infection (7,8).

Mice harbouring the MHCI allele H-2L^d^ (e.g. BALB/c) control *Toxoplasma* infection due to an immunodominant epitope derived from the GRA6 parasite protein (9–11). BALB/c mice exhibit very few tissue brain cysts and the functionality of their CD8 T cells in the *Toxoplasma-infected* brain is defined by their capacity to produce IFNγ and perforin (7,12,13). Recently, using the murine BALB/c chronic *Toxoplasma* model, a T cell population (T_int_) in an intermediate state between effector and memory status was discovered, highlighting the value of this model for defining the fate of CD8 T cells during chronic infection with persistent antigen (14).

In addition to memory T-cell populations, distinct memory T-cell population that persist long term within non-lymphoid tissue have recently been documented and are resident in nature, self-renewing, and highly protective against subsequent infections (15,16). These are termed resident memory T cells (T_RM_) and can be identified by CD103 expression (17,18). Most T_RM_ cells to date have been characterised in mucosal tissue sites, where they are rapidly active against secondary infections (19–21). Much less is known about T_RM_ in the CNS. Viral models have defined CD8 T_RM_ in VSV encephalitis and in inoculation with LCMV (15,20–22). In a susceptible C57BL/6 model of *Toxoplasma* infection, a transcriptionally defined resident memory CD8 population was recently defined in the brain (23). Again, prerequisites in terms of TCR-MHC affinity for the transition of CD8 T cells to a TRM phenotype are completely unexplored.

Rather than varying the antigenic peptide, we sought to use distinct clonal T cells. In order to answer how TCR-MHC affinity dictates trafficking and phenotype of memory CD8 T cells in the brain during chronic infection, we employed three distinct clonal CD8 T cells, each expressing a natural TCR recognizing the *Toxoplasma* antigen ROP7 (24,25). These cells were obtained from transnuclear (TN) mice generated by somatic cell nuclear transfer from a nucleus of a *Toxoplasma* antigen-specific CD8 T cell and have different affinity for MHC class I loaded with the same ROP7 peptide (24,25).

Here we report that TCR-MHC affinity dictates the potential of a CD8 T cells to home to the *Toxoplasma-infected* brain. We employed three natural CD8 T cell clones derived from a resolving *Toxoplasma* infection by somatic cell nuclear transfer, defined to possess different affinities for the same *Toxoplasma* antigen ROP7 (24,25). The two T cell clones with higher affinity, R7-I and R7-III were found in the brain during chronic infection, while the lowest affinity clone R7-II was not, despite all three clones being activated during the acute phase of infection. As possible causes for this divergent homing we observed high expression of the negative regulator of T cell activation Klf2 and its regulated genes in peptide-activated R7-II T cells. Additionally, Ctla4, a negative regulator of T cell responses was also upregulated on R7-II T cells. The highest affinity clone, R7-I, persisted longer during the chronic phase of infection than R7-III and was able to generate more T_RM_ cells in the brain. Thus, our results indicate that higher affinity of the TCR-MHC interaction is better for trafficking and persisting of the specific CD8 T cells at the site of chronic infection, here brain.

## Materials and Methods

### Mice

Thy1.1 (BALB/c N4; CD90.1^+^) and transnuclear (TN) R7-I, -II and -III mice on a Rag2 proficient BALB/c (Rag2^+/+^CD90.2^+^) background were housed and bred in the animal facility of the Francis Crick Institute (Mill Hill Laboratory, London, UK) (24). All experiments were performed in accordance with the Animals (Scientific Procedures) Act 1986.

### Calcium flux assay

For calcium flux measurements lymphocytes from lymph nodes of R7-I, II or III mice were isolated and loaded with Indo-1 dye (Life Technologies) at concentration of 2mg/ml in IMDM media containing 5% FCS for 40 minutes at 37°C. Subsequently cells were washed 2 times with IMDM media and stained with anti-CD8 (53-6.7), anti-CD4 (GK1.5) and anti-CD3 (17A2) antibodies all from Biolegend (San Diego, CA) for 20 min at room temperature. Lymphocytes were then stimulated by addition of ROP7-MHCI dextramer (Immudex) or by addition of Ionomycin (10 ng/ml).

### ERK1/2 phosphorylation assay

For the ERK1/2 phosphorylation assay lymphocytes from lymph nodes of R7-I, II or III mice were isolated and stained with anti-CD8 (5H10, Invitrogen) and anti-CD4 (GK1.5, Biolegend) antibodies. Splenocytes loaded with ROP7 peptide were used as stimulators. Lymphocytes were then stimulated by addition of splenocytes and incubated for 0, 1, 2, 4, 8, 12 min at 37°C. At the indicated time points, cells were fixed with paraformaldehyde at a final concentration of 2%. Cells were permeabilised by addition of ice-cold 90% methanol and stored over night at −20°C. Next, cells were washed and stained with anti-pERK1/2 (pT202/pY204) (20A, BD Biosciences) and acquired using an LSR II flow cytometer. Data were analysed using Flow Jo and Prism software.

### In vitro proliferation assay

Splenocytes of R7-I, II and III mice were isolated, stained with the intracellular fluorescent dye carboxyfluorescein diacetate succinimidyl ester (CFSE; 5μΜ; Life Technologies) for 5 minutes at room temperature and plated in 96-well plates. ROP7 peptide was added in the range of concentrations from 0.5x10^−4^ to 0.5x10^−9^ M. Three days later cells were harvested and stained for FACS analysis.

### T cell adoptive transfer and infections

Lymph nodes and spleens from TN ROP7 donor mice were harvested and the released cells negatively selected for CD8 T cells. Recipient Thy1.1 (BALB/c) mice received 10^6^ ROP7^+^ CD8 T cells via i.v. injection prior to infection. Mice were infected orally with 5 cysts of the ME49 *Toxoplasma* strain. Cells were harvested at the indicated time points during the acute and chronic phases of infection and processed accordingly.

### Isolation of brain mononuclear cells

Isolation of brain mononuclear cells was performed as described before (26). Briefly, mice were perfused with cold PBS and brains were isolated and homogenised. Brain cell suspension was diluted to 30% with isotonic Percoll solution and layered on top of 70% isotonic Percoll solution. Gradients were spun for 30 min at 500 x g, 18°C. Mononuclear cell population was collected from the interphase of Percoll gradient, washed and resuspended for antibody staining or restimulation.

### In vivo proliferation assay

ROP7 CD8 T cells were prepared as described. Prior to subcutaneous injection into recipient mice, cells were stained with CFSE (5μΜ). Mice were then infected orally as described. Spleen, LN and mLN tissues were then harvested 6/7 days post-infection and processed accordingly.

### Ex vivo functional assay

Cells harvested from mice 3 weeks post infection (weeks p.i.) were cultured *ex vivo* as a cell suspension for 2 hours with Ionomycin (20 ng/ml) and PMA (1 μg/ml) after which Brefeldin A (2 μg/ml) was added for the next 2 hours in RPMI media. Cells were then stained for flow cytometry and analysed as described.

### Micropipette 2-dimentional adhesion frequency assay

The two dimensional affinities were measured by micropipette adhesion frequency assay (27). CD8 T cells were negatively selected by magnetic cell sorting (Milteney) from spleen and lymph nodes of TN ROP7 mice. Human red blood cells RBCs were isolated in accordance with the Institutional Review Board at Emory University and used as the surrogate APC sensor through incorporation of ROP7 monomers with mouse b_2_-microglobulin (National Institutes of Health Tetramer Core) via biotin:strapavidin interactions. RBCs were coated with Biotin-X-NHS (EMD, Millipore) and 0.5 mg/ml streptavidin (Thermo Fisher Scientific) and 1-2 mg of the monomer antigenic and control monomers. Monoclonal cells were brought into contact 50 times with pMHC coated RBC’s with the same contact time and area (A_c_), and the adhesion frequency (P_a_) was calculated. Quantification of binding events along with TCR and p-MHC surface densities and adhesion frequencies along with two dimensional affinity were as described (27,28).

### Antibodies

Fluorescently labelled antibodies against CD3, CD90.1, CD90.2, CD62L, CD127, CD103, KLRG1, PD1 antigens and IFNγ were purchased from Biolegend (San Diego, US). Fluorescently labelled antibodies against CD8 (5H10) alpha and anti-CD69 were purchased from Life Technologies (Carlsbad, US). H-2L^d^ monomers with IPANAGRFF or photo-cleavable peptide (YPNVNI(Apn)NF) were obtained from the NIH Tetramer Core Facility (Emory University, Atlanta, US) and were tetramerised and peptide-exchanged as described previously {Frickel:2008ex}[50]{Frickel:2008ex}. All peptides were synthesised by Pepceuticals (Leicestershire, UK).

### Flow cytometry

Single-cell suspensions were prepared from brain, spleen, lymph nodes (LN) and mesenteric LN (mLN) by mechanical disruption. Brain mononuclear cells were isolated as described above (26). Cell were stained for 20 min at 4°C in an appropriate antibody cocktail and washed with PBS with 1% BSA. BD Cytofix/Cytoperm kit was used for intracellular staining of cells. Cells were run on a BD LSRII or BD Fortessa X20 and analysed using FlowJo software (Tree Star).

### Chimeras

Recipient BALB/c mice (CD90.1) were treated with an intraperitoneal injection of myeloablative agent Busulfan (10 mg/kg) and injected with a congenic (CD90.2) donor bone marrow (BM) from ROP7 transnuclear mice one day after to create bone marrow chimeras. 6 to 8 weeks after BM transplantation chimerism was assessed in the blood and mice were infected orally with 5 cysts of *Toxoplasma* ME49.

### RNASeq analysis

Single-cell suspensions of splenocytes from R7-I, -II or -III mice were incubated in RPMI medium 1640 supplemented with recombinant mouse IL-2 (10 ng/ml) overnight at 37°C with or without ROP7 peptide (10 μΜ). Cells were stained with live/dead, anti-CD3, anti-CD8a and ROP7 tetramer. Live CD8^+^ tetramer T cells were sorted on the Aria, XDP and Influx 1 cell sorters. Samples were maintained at 4°C and purity determined to be 90-95%. RNA was isolated using Trizol and the RNeasy Micro-Kit (Qiagen). A total of 200 ng of RNA was used to prepare the RNA library using TruSeq mRNA Library Prep Kit v2 (Illumina) according to the manufacturer’s recommendations. RNA sequencing was performed on the Illumina HiSeq 2500 and typically generated ~25 million 100bp non-strand-specific single-end reads per sample. The RSEM package (version 1.2.11) (29) was used for the alignment and subsequent gene-level counting of the sequenced reads relative to mm10 RefSeq genes downloaded from the UCSC Table Browser (30) on 27^th^ May 2015. Differential expression analysis between the triplicate groups was performed with DESeq2 (version 1.8.1) (31) after removal of genes with a maximum transcript per million (TPM) value of 1 across all samples in the experiment. Significant expression differences were identified at an FDR threshold of 0.01. Gene set enrichment analysis was performed by Gene Ontology Biological processes using GeneGo MetaCore (https://portal.genego.com/). Pathway analysis was performed using IPA software to demonstrate the biological effect of differentially expressed genes on cell cycle progression.

### Real-Time PCR

RNA was extracted from ROP7-specific splenocytes, either straight from the spleen, or incubated overnight with or without ROP7 peptide using RNeasy Mini and Micro Kits (Qiagen). cDNA was synthesised using the Maxima first strand cDNA synthesis kit for RT-qPCR, with dsDNase (Thermo Scientific).

Quantitative real-time PCR was performed using Maxima SYBR Green/Rox qPCR master mix (Thermo Scientific).

Results were normalised to the expression of CD8. Relative fold change was calculated by normalising to the average of R7-I, R7-II or R7-III biological triplicates respectively (straight from the spleen)

Primers used were as follows: KLF2 forward 5’ - TGTGAGAAATGCCTTTGAGTTTACTG-3’, reverse 5’- CCCTTATAGAAATACAATCGGTCATAGTC- 3’, CXCR3 forward 5’- GCCAAGCCATGTACCTTGAG- 3’, reverse 5’- GTCAGAGAAGTCGCTCTCG- 3’, Sell forward 5’- ACGGGCCCCAGT GT CAGT AT GT G-3’, reverse 5’-GATGCTGTTGTTGTACTTCGGG-3’, reverse 5’- ACCACTGAGGCATTAGAGAGC- 3’,CXCR4 forward 5’- GACTGGCAT AGTCGGCAAT G-3’, reverse 5’- AGAAGGGGAGT GT GAT GACAAA-3’,IL-7Rα forward 5’-GCGGACGATCACTCCTTCTG-3’, reverse 5’- GCATTTCACTCGTAAAAGAGCCC-3’, CD8 forward 5’- GATATAAATCTCCTGTCTGCCCATC-3’, reverse 5’- ATTCATACCACTTGCTTCCTTGC-3’.

### Statistical analyses

GraphPad software (Prism) was used to perform statistical tests. Comparisons between two groups were made using Student’s t test. Comparisons between multiple groups were made using one-way analysis of variance (ANOVA) test. Levels of significance are denoted as follows: * p ≤ 0.05, ** p ≤ 0.01, *** p≤ 0.001, **** p ≤ 0.0001. Non-significant results are either not marked or indicated as NS.

## Results

### Three CD8 T cell clones specific for the same peptide respond differently to in vitro TCR stimulation with cognate antigen

We previously described transnuclear mouse lines (24), which we used herein as a source of three CD8 T cell clones specific for the same peptide (IPAAAGRFF) derived from the ROP7 protein of *Toxoplasma gondii.* We refer to these CD8 T cell clones as R7-I, R7-II and R7-III CD8 T cells (24). We previously showed these CD8 T cell clones to differ in their TCR 3D affinity to cognate ROP7 peptide, with R7-I being the strongest binder at 4 μΜ and R7-II the weakest at 109 μΜ. R7-III has a binding affinity of 24 μΜ (25). To further define the kinetics of TCR signalling after stimulation with ROP7 peptide, we measured ER-driven calcium release, phosphorylation of ERK1/2 kinase and cell proliferation as determinants of TCR reactivity. Both the calcium release and phosphorylation assays reflected the hierarchy of the TCR-MHC binding 3D affinity and were the fastest and strongest in R7-I CD8 T cells, while the R7-II CD8 T cell response was lowest (Fig 1A and B). Additionally, we noted that R7-II CD8 T cells had a basal level of free intracellular calcium that was higher than that of R7-I and R7-III CD8 T cells (Fig 1A). In the *in vitro* proliferation assay R7-II CD8 T cells were not able to proliferate efficiently even at the highest (500 μΜ) concentration of ROP7 peptide loaded onto splenocytes while R7-I and R7-III CD8 T cells reached the highest division index at concentrations 5 μΜ and 0.5 μΜ respectively (Fig. 1C). These *in vitro* experiments suggest that the 3D surface plasmon resonance affinity of the TCR-MHC binding reflects the strength of downstream signalling and partially translates to proliferation capacity *in vitro*.

**Figure 1.**
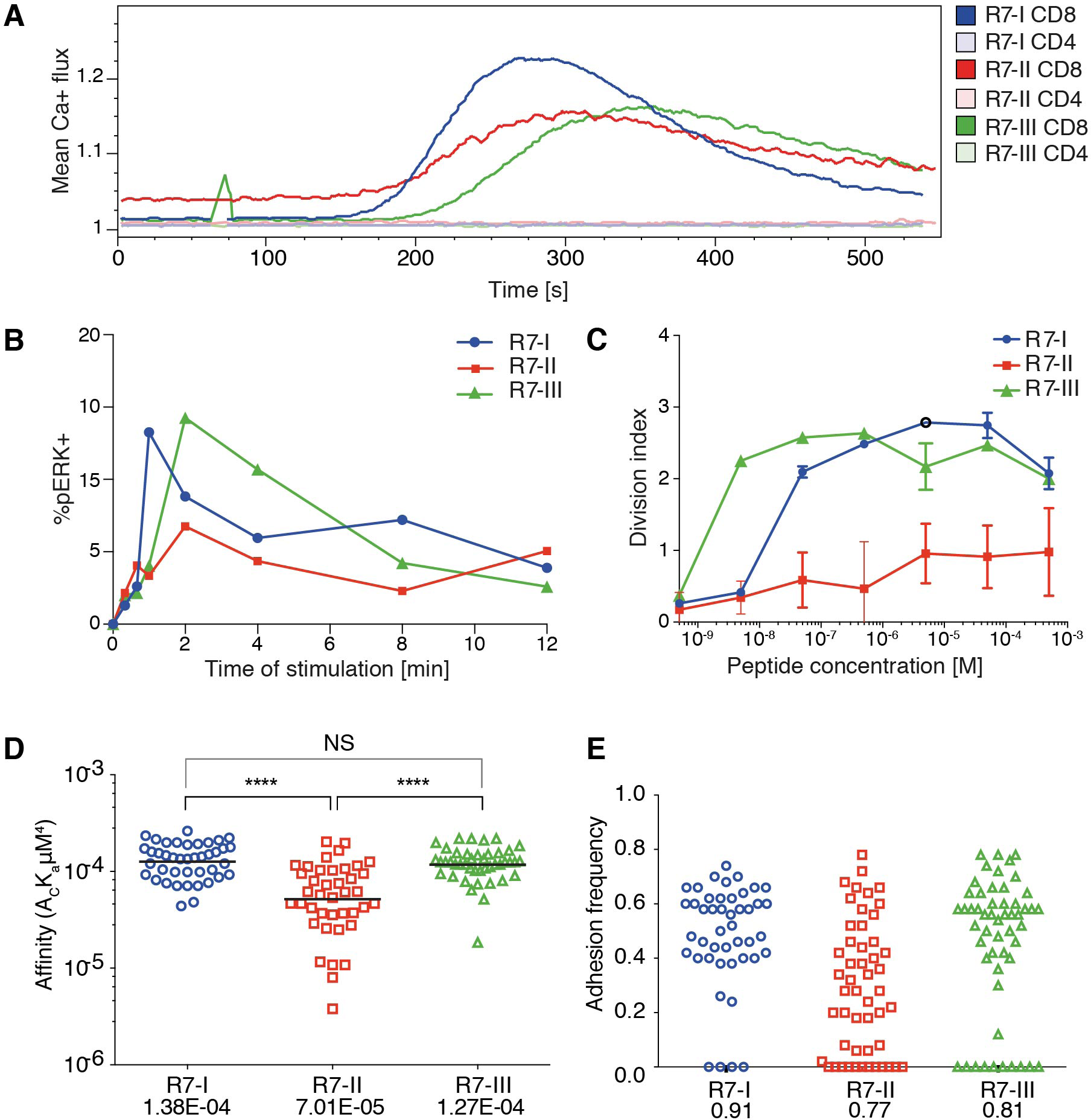
Three distinct CD8 T cell clones of the same specificity exhibit different 2D affinity and responses to specific TCR receptor stimulation. A) Ca^2+^ flux profile of the three R7 CD8 T cell lines upon stimulation of TCR signalling with ROP7 dextramer (representative of two experiments with multiple mice each). B) Phosphorylation of ERK1/2 kinase in time upon stimulation with splenocytes loaded with ROP7 peptide (representative of two experiments). C) Proliferation of R7 CD8 T cells after stimulation with a range of ROP7 peptide concentrations (representative of three experiments). D) Relative 2D affinity of the three R7 CD8 T cell clones specific for ROP7 (IPAAAGRFF) normalized by TCR surface density. Each individual data point represents the affinity for a single CD8 T cell. Analysis utilizes the geometric mean of the population. E) Adhesion frequencies of the three ROP7 clones F) Gaussian curves were fitted to the affinities of (D) Blue line represents R7-I, red line represents R7-II and the green line represents R7-III. Data are cumulative from two individual experiments and a total of 3 mice per clone.

To provide additional insight into the functional response of the three R7 CD8 T cell clones during *Toxoplasma* infection, we determined their 2D affinity for the ROP7 antigen. The micropipette adhesion frequency assay provides 2D based measures of TCR affinity for pMHC in a context that is membrane anchored. 2D affinity correlate more closely to functional responses than do 3D affinity measurements whose measurements are based on purified proteins. Over 40 T cells for each clone were analysed to reveal similar high affinities for R7- and R7-III (geometric means being 1.38E-04 and 1.27E-04 respectively) and that R7-I whilst having a similar affinity to R7-III (Fig 1D) has a higher adhesion frequency than R7-III (Fig 1E) that being 0.91 and 0.81 respectively. R7-II had a 3-fold lower 2D affinity (7.01E-05). The 3-fold difference in affinity is functionally relevant as previously, we have demonstrated that during Polyoma infection CD8 T cells with the highest 2D affinity are found in the CNS and eventually comprise the T_RM_ population (32). In addition, we have reported CD4 T cells mediating EAE carry a 2-fold higher affinity as compared to the peripheral T cells (33).

### All three clones of R7 CD8 T cells are efficiently primed in the acute phase of Toxoplasma infection

Next, we sought to verify if these differences observed *in vitro* would still hold true *in vivo.* R7 CD8 T cells were adoptively transferred into congenic naïve recipient mice (CD90.1 BALB/c). Subsequently, mice were orally infected with ME49 *Toxoplasma* tissue cysts. The donor R7 CD8 T cells could then be followed during the acute and chronic phase of infection (Fig. 2A). We were able to observe proliferated cells for all three R7 clones in the mLN earliest 6 days post infection (p.i.) (Fig. 2B). Limited number of R7-II donor cells could be found in the mLN. However, those that were recovered from mLN had low CFSE level indicating that they had proliferated similarly to R7-I and R7-III CD8 T cells. Additionally, the R7 donor CD8 T cells were all activated to the same extent based on CD69 expression (Fig. 2B). We conclude that all three R7 clones are responsive to a *Toxoplasma* infection *in vivo*, as measured by proliferation and activation status.

**Figure 2.**
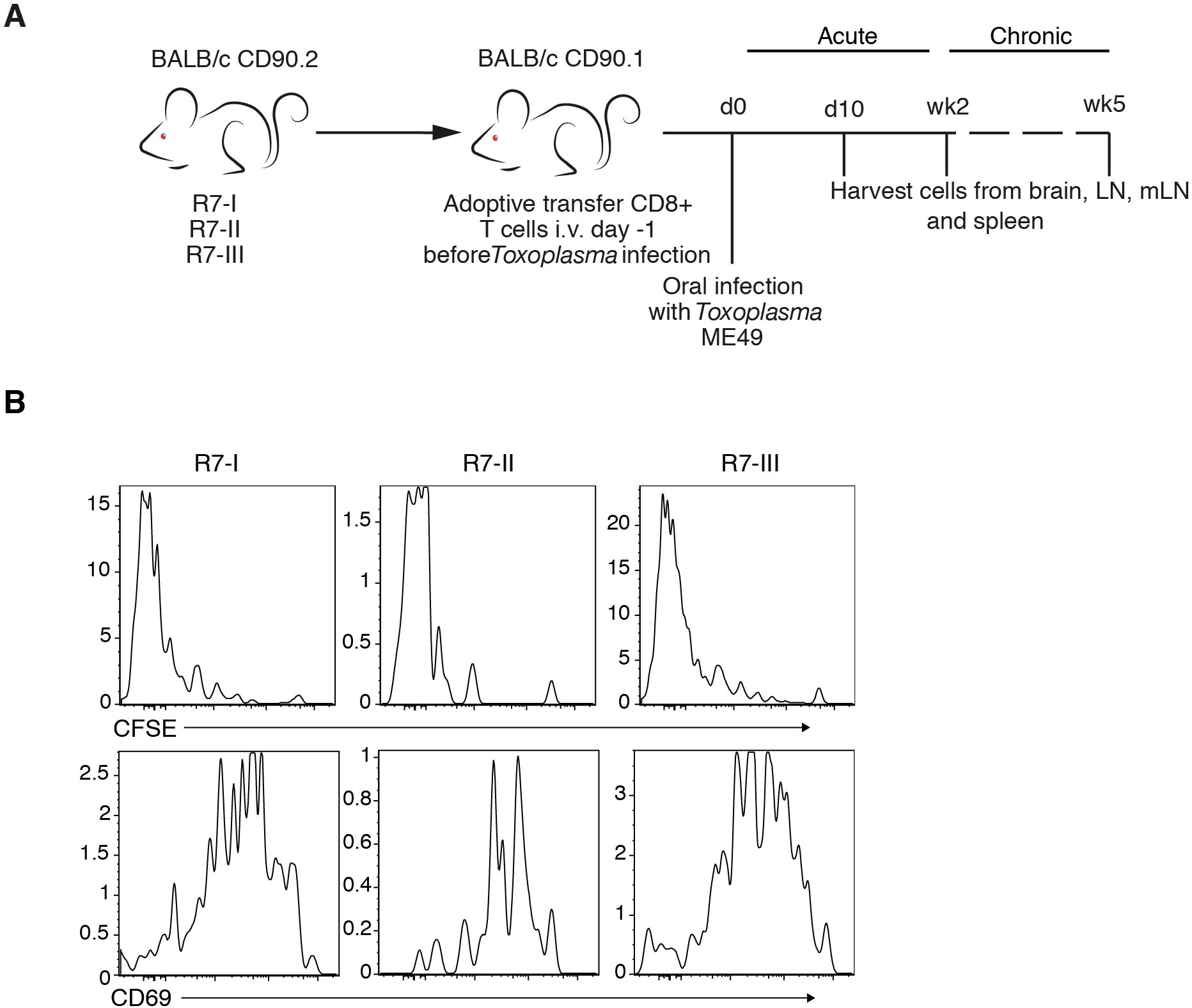
Three clones of R7-reactive CD8 T cells are primed and activated during the acute phase of Toxoplasma infection. A) Schematic diagram of the experimental set-up for *in vivo* experiments. Transnuclear R7 mice were used as a donor of CD8 T cells in adoptive transfer experiments (AT). CD8 T cells isolated from lymph nodes and spleens were transferred to congenic recipients. Recipient mice were infected orally with 5 cysts of *Toxoplasma* ME49. At different time points post-infection (p.i.) recipient mice were sacrificed and donor CD8 T cell populations were assessed. B) Donor cells were stained with CFSE before transfer. Histograms show CFSE dilution and CD69 expression on transferred R7 CD8 T cells isolated from mesenteric (mLN) 6 days p.i. (representative of two independent experiments).

### R7-II CD8 T cells do not persist and do not reach the brain of recipient mice during Toxoplasma infection

Cysts in the brain characterise the chronic phase of *Toxoplasma* infection. IFNγ produced by CD8 T cells is crucial for the maintenance of the quiescent cyst form of *Toxoplasma* (8,13). We showed that the three R7 CD8 T cell clones could be primed and proliferated in the acute phase of *Toxoplasma* infection. Next, we investigated if R7 CD8 T cells could be found in the brain in the chronic phase of *Toxoplasma* infection. We analysed brain, spleen, mLN and nondraining LN for the presence of transferred R7 CD8 T cells 3 weeks p.i̤ R7-I and R7-III CD8 T cells were found in significant numbers in the brain at 3 weeks p.i. (Fig. 3A). Percentages and absolute numbers of donor R7 CD8 T cells of all CD8 T cells in a given organ were different depending on the clone (Fig. 3A). The R7-II clone was found in insignificant percentages and numbers in all tested organs. R7-I and R7-III clones were present in higher percentages from 2-15% of all CD8 T cells depending on the organ. There was no significant difference between percentages of the R7-I and R7-III clone in the brain. In the spleen and mLN we observed significantly higher percentages of the R7-III CD8 T cells. We also determined absolute cell number for each clone at each site and did not observe significant differences between the R7-I and R7-III clone (Fig. 3A right panel).

**Figure 3.**
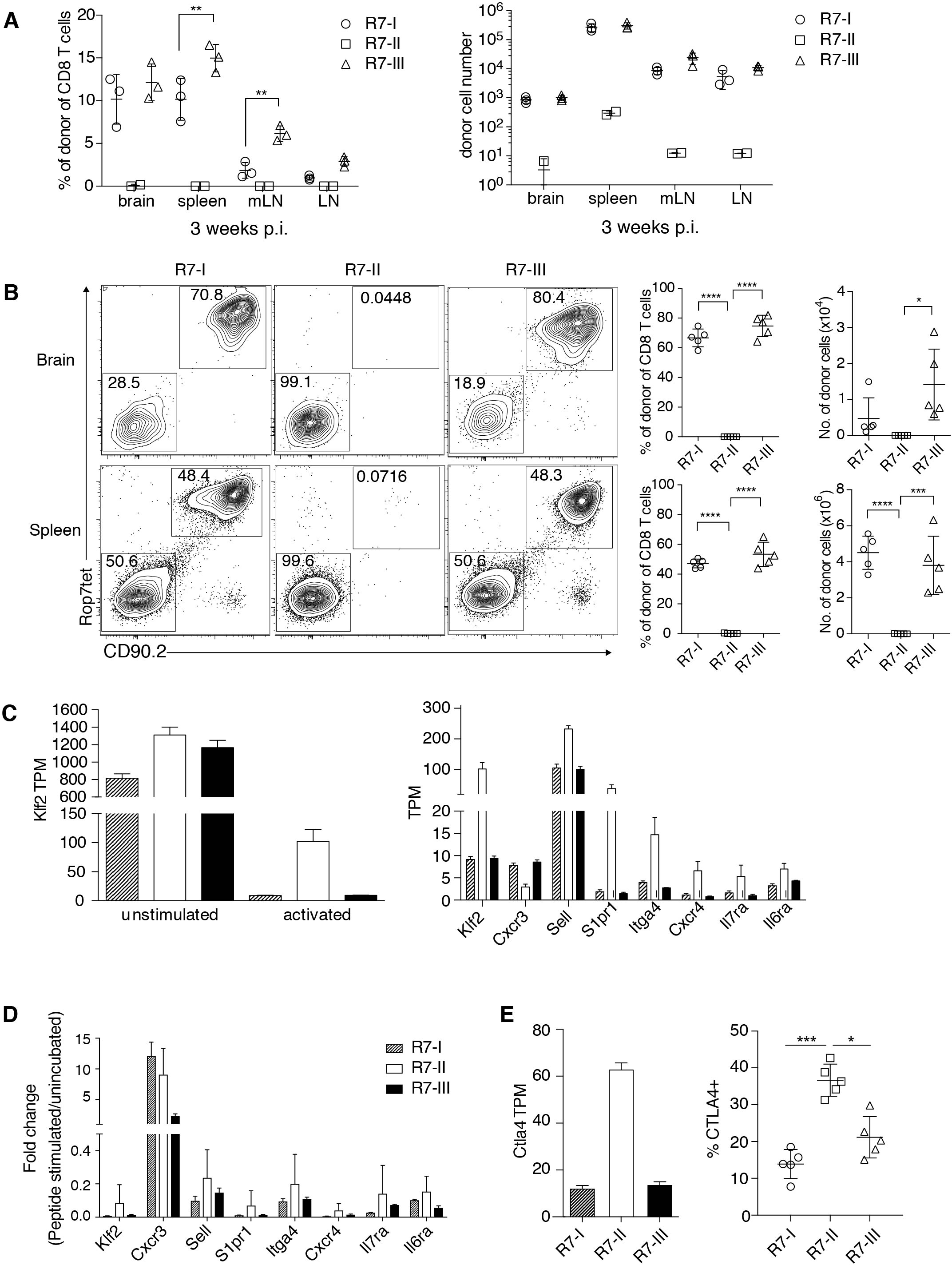
R7-II CD8 T cells do not persist and do not reach the brain of recipient mice during *Toxoplasma* infection. A) Percentages and total cell numbers of donor ROP7 CD8 T cells in the brain and lymphoid organs 3 weeks p.i. (representative of at least 5 experiments with 3 mice per line), * p ≤ 0.05, ** p ≤ 0.01, *** p ≤ 0.001, 2-way Anova followed by multiple comparisons Tuckey’s. Mean and SD. B) Percentages and cell numbers of donor R7 CD8 T cells in the brain and spleen in the acute phase of infection (day 10 p.i.) (representative of two experiments), * p ≤ 0.05, ** p ≤0.01, *** p ≤ 0.001one-way Anova followed by multiple comparisons Tuckey’s. Mean and SD. C) Splenocytes of R7 mice were left untreated (unstimulated) or stimulated with the Rop7 peptide (activated) over night. On the next day ROP7tet^+^ CD8 T cells were sorted and lysed for RNA extraction. Transcripts levels were evaluated in an RNAseq experiment. Expression of *Klf2* and Klf2-regulated genes is shown as a number of transcripts per million (TMP) in unstimulated and activated (left hand graph) or only activated R7 CD8 T cells (right graph). D) Validation of RNAseq results was performed on samples from an independent experiment with use of qRT-PCR. Expression of *Klf2* and Klf2-regulated genes shown as normalized 2^^^- ΔΔCt; values. E) *CTLA4* expression in the RNAseq experiment shown as number of TMPs in activated samples (left graph) and CTLA4 protein expression evaluated by FACS of cells isolated 5 days p.i. from LN of mice adoptively transferred with R7 cells and infected with *Toxoplasma* (right graph). * p ≤ 0.05, ** p ≤ 0.01, *** p ≤ 0.001.

We analysed earlier time points to estimate the time when donor cells reached the brain and to assess if there is a difference in donor cell number or percentages in the acute phase of infection. R7-I and R7-III CD8 T cells were observed in the brain as early as 10 days p.i., however, we failed to detect a distinct R7-II CD8 T cell population in the brain (Fig. 3B). R7-I CD8 T cells could not be observed in prominent numbers in any of the tested primary and secondary lymphoid organs suggesting a lack of proper expansion and homing of the transferred population to the brain (Fig. 3B and Fig. S1A). R7-I and R7-III CD8 T cells were a major part (60-80%) of the total CD8 T cell population in the brain at day 10 p.i̤ Their percentages decreased throughout time while host CD8 T cells reached the brain suggesting that the transferred T cell clones had a head start compared to newly formed *Toxoplasma*-specific CD8 T cells of the host. There were no significant differences in percentages or numbers of donor population between R7-I and R7-III CD8 T cells at day 10 or 2 weeks p.i. (Fig. 3B and Fig. S1A).

To dissect the reasons for the poor expansion and lack of R7-II cells in the brain we compared the transcriptional profiles of *in vitro* ROP7-activated R7-II vs R7-I or R7-III cells. In the list of the top 10 genes upregulated in R7-II (Table 1) we found *Klf2* encoding a transcription factor that is known to be important for T cell trafficking between the blood and lymphoid organs (34–36). Klf2 is highly expressed in naïve and memory T cells but downregulated in effector T cells upon binding of the TCR to its cognate peptide (37). The stronger the binding affinity, the lower the expression of *Klf2* and the better the activation of the T cells (38). Additionally, *S1pr1* which is regulated by Klf2 was also in the top 10 upregulated genes. We analysed the expression of *Klf2* across the unstimulated and activated samples of the RNASeq experiment (Fig. 3C, left graph) as well as at the genes known to be regulated by Klf2 such as *CXCR3, Sell (CD62L), S1pr1, Itga4, CXCR4, Il7ra, Il6ra* (Fig. 3C, right graph). *Klf2* expression was at similar level in all three naïve CD8 T cell clones. After activation, all three clones downregulated *Klf2* however downregulation in R7-II was the weakest reflecting the lowest binding affinity of Rop7 peptide to R7-II. Expression of *Sell (CD62L), S1pr1, Itga4, CXCR4, Il7ra, Il6ra* mirrored *Klf2* expression being highest in R7-II and was comparable between R7-I and R7-III. On the other hand, *CXCR3* expression was lowest in R7-II. We performed an independent *in vitro* activation experiment and confirmed by qRT-PCR the RNAseq results observed (Fig. 3D). mRNA expression detected by qRT-PCR correlated with the RNAseq data for all tested genes besides the *CXCR3* gene.

**Table 1.** Top 10 up and downregulated genes in activated R7-II vs R7-I and R7-III. Table shows top 10 up and downregulated genes in R7-II samples as compared with R7-I and R7-III samples upon activation, where only genes differentially expressed in R7-II vs R7-I and R7-II vs R7-III comparisons but similar expression between R7-I and R7-III were included. For each gene in the table fold change (FC) and false discovery rate (FDR) is shown form R7-I vs R7-III comparison.

|  | Gene symbol | FC | FDR |
| --- | --- | --- | --- |
| 1 | Tnfrsf19/CD137 | 19.20 | 5.84E-105 |
| 2 | Mturn | 14.93 | 1.95E-77 |
| 3 | Rasgrp2 | 13.49 | 1.27E-90 |
| 4 | Slc6a19 | 12.84 | 2.18E-44 |
| 5 | Cd7 | 11.38 | 4.03E-49 |
| 6 | S1pr1 | 9.77 | 1.21E-44 |
| 7 | Nsg2 | 9.68 | 3.28E-34 |
| 8 | Klf2 | 8.80 | 4.88E-149 |
| 9 | Atp6v0e2 | 8.60 | 1.18E-29 |
| 10 | Arl4c | 8.45 | 5.70E-105 |

**Table 1. Top 10 up and downregulated genes in activated R7-II vs R7-I and R7-III**
|  | Gene symbol | FC | FDR |
| --- | --- | --- | --- |
| 1 | Tbx21/Tbet | -6.45 | 4.64E-38 |
| 2 | Tnfsf11/RANKL | -6.07 | 5.80E-160 |
| 3 | Serpinb6b | -5.64 | 2.11E-42 |
| 4 | Dapl1 | -5.51 | 1.15E-38 |
| 5 | Ccl9 | -5.46 | 3.81E-93 |
| 6 | Rnf157 | -4.89 | 5.80E-78 |
| 7 | Tbkbp1 | -4.51 | 2.74E-32 |
| 8 | Lipg | -4.51 | 4.47E-72 |
| 9 | Stc2 | -4.14 | 8.38E-16 |
| 10 | Chst11 | -3.95 | 1.98E-56 |

CTLA4 is known as a negative regulator of T cell responses. We analysed the expression of the *Ctla4* transcript in the RNASeq of activated ROP7 CD8 T cells as its high expression in R7-II cells could explain their poor performance. As expected, we observed the highest expression of *Ctla4* in R7-II CD8 T cells in comparison with the two other R7 clones (Fig. 3E, left graph). We could confirm the RNAseq data in an *in vivo* experiment, where R7 T cells were adoptively transferred to congenic mice and analysed in LNs 5 days after infection with *Toxoplasma*. CTLA4 surface expression determined by FACS in R7-II cells was highest in comparison to R7-I and R7-III CD8 T cell clones (Fig. 3D, right graph).

### R7-III has a higher contraction rate and does not persist in the late phase of chronic infection

At 5 weeks p.i. compared to 3 weeks p.i., we observed a dramatic decrease in the R7-III CD8 T cell population in all of tested lymphoid and non-lymphoid organs while the R7-I CD8 T cell population decreased more subtly, which could be explained by natural contraction of the population after an initial expansion phase (Fig. 4A and Fig. S1B).

**Figure 4.**
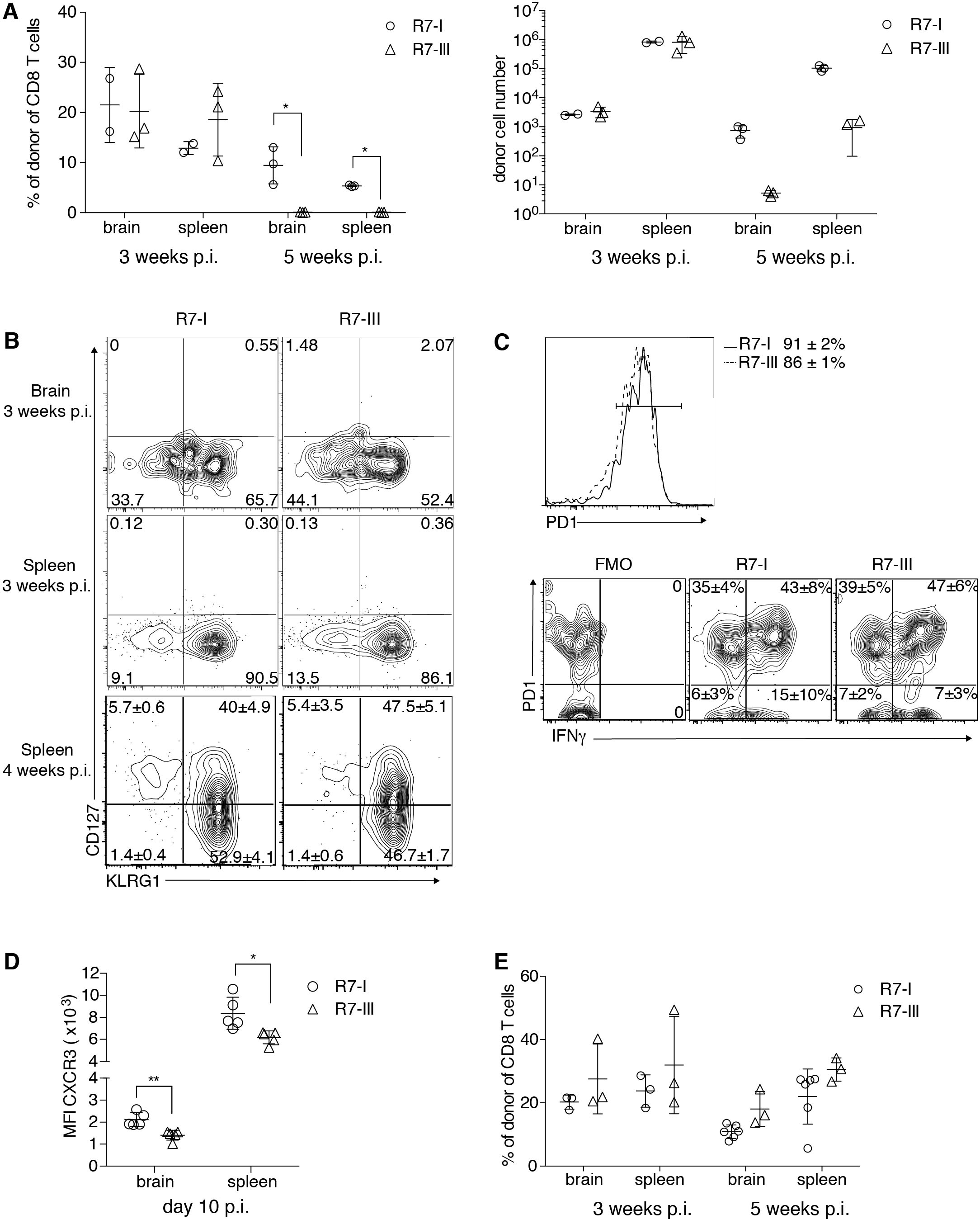
R7-III has a higher contraction rate and does not persist into the late phase of the chronic infection. ROP7 CD8 T cells were adoptively transferred into congenic mice before *Toxoplasma* infection. Spleen, mesenteric LN and popliteal and axillary LN and brains were harvested and analysed by flow cytometry (A-E). A) Percentages and total cell numbers of donor Rop7 CD8 T cells in the brain and spleen 3 and 5 weeks p.i. (representative of at least 5 experiments). * p 0.05, ** p ≤ 0.01, *** p ≤ 0.001, Two-way Anova followed by Tuckey multiple comparisons test. B) Donor population was assessed for expression of CD127 (IL7Ra) and KLRG1 at 3 and weeks p.i̤ (representative of at least 5 experiments) C) PD1 expression on donor CD8 T cells in the brain 3 weeks p.i. and their potential to produce IFNγ after *ex-vivo* restimulation with PMA/Ionomycin, (representative of 2 experiments). * p ≤ 0.05, ** p ≤ 0.01, *** p ≤ 0.001 by Student t-test. D) CXCR3 surface staining on donor CD8 T cells 10 days p.i. in brain and spleen. E) Percentages of R7 CD8 T cells in brain and spleen of R7 bone marrow chimeras at and 5 weeks (minimum of 3 mice analysed per condition) p.i̤ * p ≤ 0.05, ** p ≤ 0.01, *** p 0.001 by Student t-test.

The formation of memory CD8 T cells in persistent infections is still controversial (39). In BALB/c mice where persistent *Toxoplasma* infection is known to be controlled by CD8 T cells (13), the presence of memory CD8 T cells is expected. We analysed if R7 CD8 T cells differentiated into memory cells and if the ratio of differentiation into effector and memory precursors was the same between R7-I and R7-III CD8 T cells. Short lived effector cells (SLEC, CD127^-^KLRG1^+^) were present in similar percentages in R7-I and R7-III CD8 T cell populations in the brain and spleen at 3 and 4 weeks p.i. (Fig. 4B). Memory precursor effector cells (MPEC, CD127^+^KLRG1^-^) where not present at 3 weeks p.i., however we detected population of MPEC at 4 weeks p.i. in the spleen in similar percentages between R7-I and R7-I CD8 T cell (Fig. 4B, bottom panel).

In the chronic phase of *Toxoplasma* infection in C57BL/6 mice CD8 T cells in the brain are exhausted and express high levels of the exhaustion marker PD1 (40). Blockade of the PD1–PDL1 pathway has been shown to rescue the exhaustion phenotype of CD8 T cells and prevent mortality of chronically *Toxoplasma* infected animals (41). In contrast to C57BL/6 mice, BALB/c mice are resistant to chronic *Toxoplasma* infection (9) and *Toxoplasma* GRA6-specific CD8 T cells in BALB/c mice lack PD1 expression during the chronic phase (14). We investigated if R7 CD8 T cells in brain of BALB/c mice at 3 weeks p.i. were exhausted. Almost 90% of R7 CD8 T cells expressed PD1 (Fig. 4C). No significant difference was observed between R7-I and R7-III CD8 T cell populations in the brain suggesting that it is not exhaustion that leads to greater contraction of the R7-III CD8 T cell population. As PD1 can be also a marker for recently activated cells we analysed the ability of PD1^+^ cells to produce IFNγ, since cytokine production is lost in a truly exhausted cell (42). Brain mononuclear cells from week of infection were *ex vivo* re-stimulated with PMA and ionomycin and stained for IFNγ (Fig. 4D). More than half of PD1 positive cells were able to produce IFNγ and only 1/3 of R7 CD8 T cells in the brain were positive for PD1 and negative for IFNγ. At the same time, only 5% of R7 cells in spleen expressed PD1 and did not produce IFNγ. No significant difference between R7-I and R7-III CD8 T cells was observed indicating that the reason for the disappearance of R7-III CD8 T cells is independent of their exhaustion state.

Next, we considered CXCR3 as a candidate molecule to unravel the differences in persistence between R7-I and R7-III. The CXCR3 receptor is important in trafficking of CD8 T cells to nonlymphoid tissues including the brain (43). We evaluated CXCR3 expression levels on R7-I and R7-III CD8 T cells during *Toxoplasma* infection. CXCR3 expression on R7-III CD8 T cells was significantly lower at day 10 p.i. both in the brain and in the spleen (Fig. 4E). This suggests that R7-III cells may have a lower ability to travel to the brain than R7-I thus amounting a difference in their presence later in infection. To better understand the observed decrease in the brain population of R7-III CD8 T cells we set up bone marrow (BM) chimeras. We used the transplant conditioning drug busulfan to induce myeloablation and create a niche for the R7-I or R7-III bone marrow (44).

In the brain and spleen, R7-I and R7-III BM chimeras have similar percentages of CD8 T cells originating from the donor’s BM at both week 3 and 5 p.i. (Fig. 4E). These results show that constant replenishment from the periphery is necessary for the persistence of donor cells is the brain at 5 weeks p.i. When R7-III CD8 T cells disappear from the periphery, they also disappear from the brain. By creating chimeras, we showed that when we keep a constant R7-III population in the periphery these cells also persist in the brain.

### R7-I CD8 T cells of T_RM_ (CD103^+^) phenotype are present in higher percentages in the brain than R7-III CD8 T cells

Previous studies have shown that tissue resident memory T cells found in the brain can survive without replenishment from the CD8 T cells circulating in the blood (15). Resident memory T cells were observed in higher percentage in R7-I than R7-III CD8 T cell populations 3 weeks p.i. in the brain (Fig. 5A). Bone marrow chimeras 3 weeks p.i. also exhibited lower percentages of R7-III T_RM_ cells in the brain (Fig. 5B) indicating that the difference to produce less T_RM_ is intrinsic to the R7-III clone independent of replenishment from periphery.

**Figure 5.**
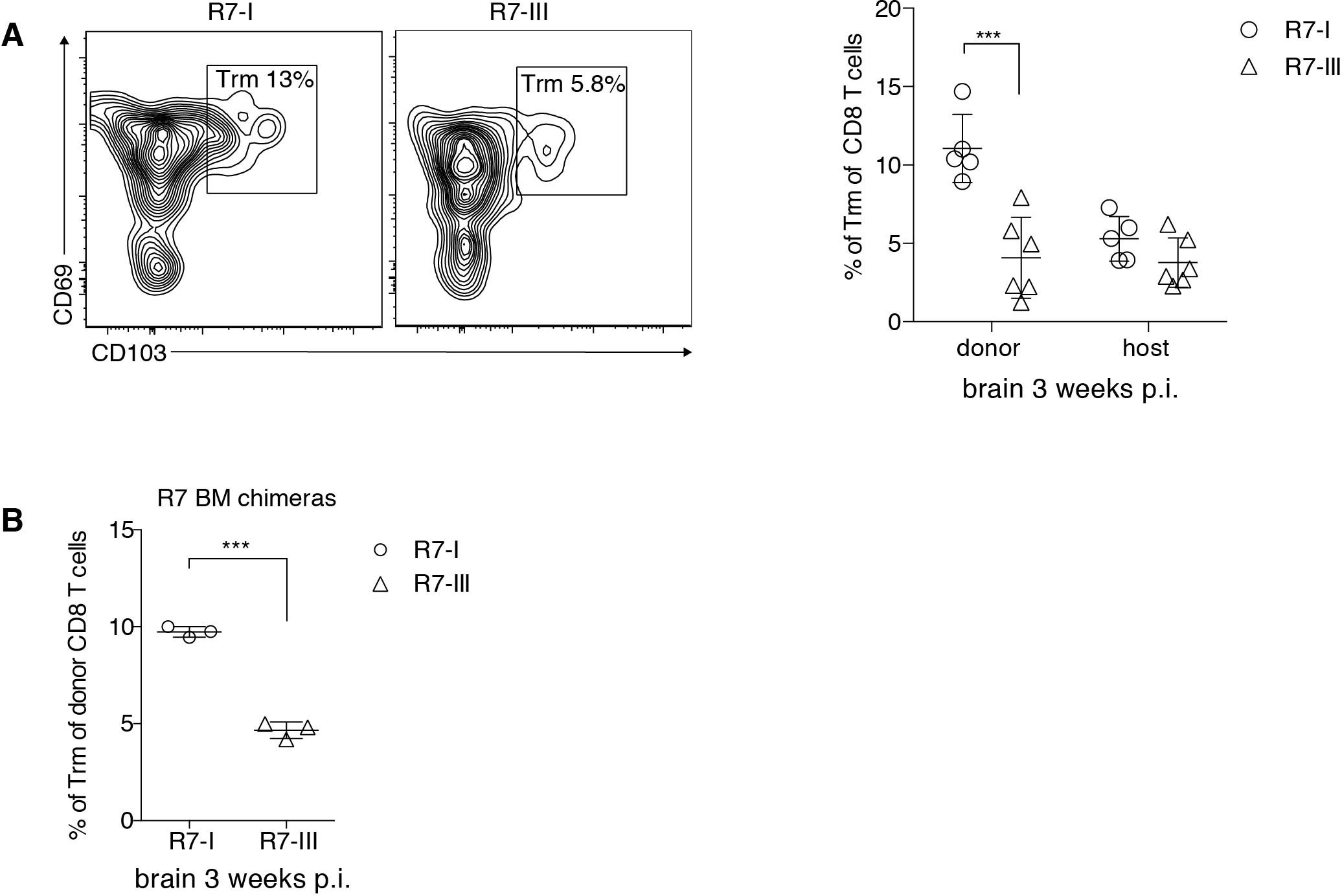
R7-I CD8 T cells of Trm (CD103^+^) phenotype are present in higher percentages in the brain than R7-III CD8 T cells. A) Representative FACS plot of donor resident memory population and percentages of donor and host CD8 T_RM_ cells in the brain 3 weeks p.i. (representative of at least 5 experiments). * p ≤0.05, ** p ≤ 0.01, *** p ≤ 0.001 2-way Anova followed by Sidak’s multiple comparisons test. B) Trm percentages in brains of R7 bone marrow chimeras mice 3 weeks p.i̤, 3 mice per T cell line, * p ≤ 0.05, ** p ≤ 0.01, *** p ≤ 0.001 by Student t-test.

## Discussion

Affinity of TCR-MHC interaction influences the fate of the activated T cell (2). Immediate effects of strong or weak interactions on activation and expansion of the T cells have been studied broadly (2,45,46). However, it is unclear how affinity influences memory CD8 T cell formation. Herein we studied three different clones of CD8 T cells (R7-I, II and III) during *Toxoplasma* infection in BALB/c mice. These three CD8 T cell clones harbour TCRs specific for the same peptide of the *Toxoplasma* protein ROP7, but differ in their sequence and affinity for that peptide presented in MHC class I (24,25). The hierarchy of affinity and functional *in vitro* responsiveness to ROP7-MHCI of the clones was R7-I>R7-III >R7-II (25). The lowest affinity clone R7-II failed to traffic to the brain during the chronic phase of infection even though we could show acute phase proliferation. R7-I outperformed R7-III in persistence in lymphoid organs and the brain in chronic infection. Additionally, R7-I was able to form more resident memory T cells in the brain than R7-III.

In order to test the affinity of the three R7 CD8 T cell clones in a more physiological setting, in addition to our previously published 3D affinity measurements, we performed 2D affinity measurements. Interestingly, we demonstrated little affinity difference between R7-I and R7-III. However, R7-I had a higher adhesion frequency than R7-III, possibly explaining the functional differences we observed between these clones in the chronic phase of infection.

R7-II, the clone with the lowest affinity for ROP7 peptide, was not found in the chronic state of *Toxoplasma* infection. Small number of the cells of that clone got activated and proliferated in the acute phase of infection, but after the contraction phase that clone could not be found neither in any analysed lymphoid organs nor in the brain. As one possible hypothesis, we demonstrated by RNAseq and subsequent qPCR that R7-II did not cross the affinity threshold required to downregulate *Klf2* levels and consequently set the cells for homing to infected tissues. Lack of homing and retention in the LN as a mechanism responsible for lack of R7-II in brain was also supported by increased CTLA-4 expression. CTLA-4 expression on T cells is known to be responsible for cells being stuck in the LN after antigen encounter and mark anergic cells (47,48). It is considered as one of T cell–intrinsic function of CTLA-4 to control self-reactive T cell motility in tissues (49). Signals from week TCR-MHC interaction may prevent full activation of the T cell, but still enable it to receive partial signals (2). Little is known about transcriptional regulation of *Ctla-4* (50). It has not been investigated if Klf2 can directly or indirectly regulate Ctla-4 expression.

R7-I and R7-III clones were of higher affinity than the R7-II clone and both were found in the brain during the chronic phase of infection. The initially quite similar clones performed differently in the later phase of infection. While both clones were able to form SLEC and MPEG memory cells, the R7-III clone did not persist in the periphery and brain in the later phase of chronic infection. Additionally, in the brain, more R7-I than R7-III cells showed a phenotype of resident memory cells (CD69^+^CD103^+^). We postulate that these differences can be attributed to the increased adhesion frequency we observed in 2D measurements, as well as the increased TCR-MHC binding affinity exhibited in 3D measurements (25). We were not able to exactly pinpoint the reason for the disappearance of R7-III cells during the chronic phase of infection. It could be attributed to slower replication, higher rate of death, or formation of different types of cells that have different abilities to survive.

R7-III has been shown to be more proliferative than R7-I (more cell cycle terms in GO analysis) (25). Also, we observed slightly higher percentages of R7-III than R7-I cells in the spleen at 3 weeks p.i. indicating that the initial slower replication rate of R7-III is not the reason for differences between R7-I and R7-III observed in the later chronic phase of infection. SLEC and MPEC percentages were not significantly different between two clones. Additionally, the expression of the exhaustion marker PD-1 and the ability to produce IFNγ also did not differ between R7-III and R7-I at 3 weeks p.i.

Creating bone marrow chimeras, we provided an artificial model where R7-specific CD8 T cells are routed from the bone marrow via the thymus to the periphery also during infection. This phenomenon has been described in persistent viral infections where host cells, but not donor cells, can be resupplied through thymic output, and new, naive specific CD8 T cells are being generated and subsequently primed during persistent infection (51). Newly generated T cells preserve antiviral CD8 T cell populations during chronic infection (51). In the C67BL/6 mice model of *Toxoplasma* infection T cells are recruited from the periphery to the brain in the chronic stage (52).

In our bone marrow chimeras model, even if cells get exhausted or/and stop dividing, new R7 specific cells are available to traffic to the brain. We concluded that constant replenishment from the periphery is necessary to keep the population of R7-III in the brain in the later stages of chronic infection. If the cells are available in the periphery they will traffic to the brain. Thus, since the R7-I clone exhibits longer survival in the periphery and in the brain without replenishment, we can conclude that cells of stronger affinity perform better in chronic infections.

The proportion of the cells with T_RM_ phenotype was different between R7-I and R7-III donor population in the brain. R7-III cells persistently exhibited a lower percentage of T_RM_ cells, no matter if able to replenish from the periphery in bone marrow chimeras or not. Additionally, R7-III cells exhibiting the phenotype of resident memory T cells (CD69^+^CD103^+^) at 3 weeks p.i. were absent from the brain at 5 weeks p.i̤ This suggests that these were possibly not true (classical) T_RM_ cells that are long lasting and shown to persist for years after infection. However, ‘classical’ T_RM_ were defined in acute infection models with rechallenge (20,53). It is thus conceivable that in chronic infection they may have a different characteristic. Constant antigen stimulation during persistent infection may have a negative influence on T_RM_ that are considered to be antigen independent. Persistent antigen stimulation has been shown to lower CD103 expression on T_RM_ but it did not block their formation (54). It is possible that these cells become exhausted when constantly stimulated and thus only newly formed CD103 positive cells contribute to the cells observed in the brain (23). It is possible that the strength of the antigen stimulation influences this process and lower affinity leads to lower number of R7-III cells expressing CD103, eventually leading to the elimination of these cells. Indeed, it has been shown that T_RM_ in the brain exhibit about 20-fold higher affinity as compared to splenic memory cells (32). As in our model R7-I and R7-III CD8 T cell are different only in their TCR receptor, we propose that observed differences are due to the affinity. One caveat is that this difference may not only be derived from the affinity of the interaction with the cognate ROP7 peptide, but different TCRs may already shape the fitness of the cells differently in the thymus (25,55). Indeed, we previously showed that R7-I and R7-III cell respond differently to CD3/CD28 stimulation with R7-III is being more proliferative. Thus, the self-reactivity of T cells may also play a role in the later stages of an infection in shaping the memory CD8 T cell phenotype.

Effector T cells in chronic infection are constantly exposed to antigen leading to exhaustion (56,57). In the *Toxoplasma* infection model of C57BL/6 mice, it has been shown that chronic infection leads to exhaustion of CD8 T cells in the brain (41,58). However, C57BL/6 mice, unlike BALB/c mice are not resistant to *Toxoplasma* and die in the chronic phase of infection (41,58). BALB/c mice, which we used in our model, are resistant to *Toxoplasma* and the chronic stage of infection is asymptomatic in this mouse strain. This implies that CD8 T cells in the BALB/c model are not exhausted. Indeed, recently published data by Chu *et al.* show that CD8 T cells specific for the immunodominant GRA6 epitope are not exhausted (14). The ROP7 epitope is subdominant, but has been shown to be well-represented in the chronic phase of infection (59). We observed high levels of PD-1 expression on brain resident R7-specific CD8 T cells. However, the majority of these cells were able to produce IFNγ and could not be perceived as exhausted. The expression of PD-1 in this case may indicate recent antigen encounter, which is not surprising in chronic infection. In contrast with what was shown for Gra6-specific cells (14), our results indicate that CD8 T cells specific for subdominant epitopes do go through a contraction phase during chronic infection. R7-I and R7-III contraction happens after a peak in T cell numbers on days 14-21.

We conclude that R7-III cells, due to their lower affinity for the ROP7 peptide form less T_RM_ cells and do not persist in the later stages of chronic infection. Our data indicate that among cells specific for the subdominant antigen the cells of higher affinity are favoured and their persistence is secured by formation of long-lasting T_RM_ cells in the non-lymphoid tissues.

## Acknowledgements

We acknowledge the NIH Tetramer Core Facility (contract HHSN272201300006C) for provision of H2Ld monomers with IPANAGRFF and photo-cleavable peptides. We would like to thank Blake Gibson for technical assistance. We are indebted to the Mill Hill Biological Services and FACS facility. We would like to thank Ben Seddon, George Kassiotis, Jean Langhorne, Charles Sinclair, Thea Hogan and Jie Yang for discussion. This work was supported by the Francis Crick Institute, which receives its core funding from Cancer Research UK (FC001076), the UK Medical Research Council (FC001076), and the Wellcome Trust (FC001076). Eva-Maria Frickel was supported by a Wellcome Trust Career Development Fellowship (091664/B/10/Z).

## Author Contributions

AS, NY, EMK conducted experiments, AS, NY, EMK, HP and BDE analysed the data, BDE provided essential technology, AS and EMF wrote the manuscript, EMF directed the study.

## Supplementary Information

**Supplementary Figure 1.**
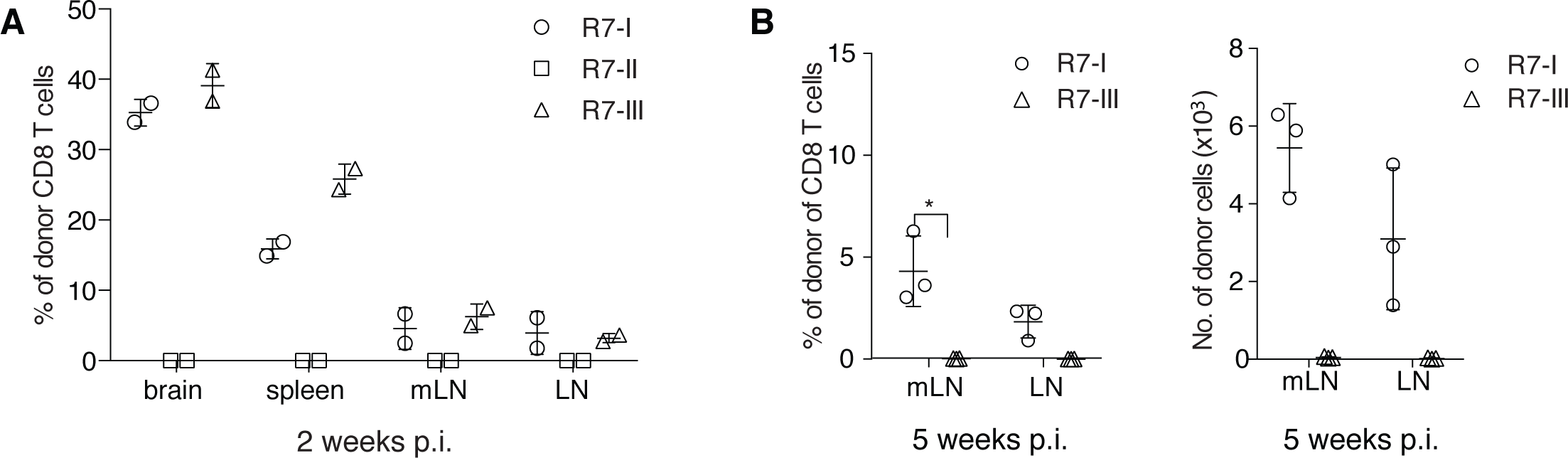
Quantification of donor R7 CD8 T cell populations at different time points post-infection. A) Percentages of donor R7 CD8 T cells in brain, spleen, mLN and non-draining LN at 2 weeks p.i. (representative of at least 5 experiments). B) Donor R7-III CD8 T cells 5 weeks p.i. in mLN and non-draining LN are present in lower percentages (left) and absolute numbers (right) than R7-I CD8 T cells (representative of at least 5 experiments).

